# A human urothelial microtissue model reveals shared colonization and survival strategies between uropathogens and asymptomatic bacteria

**DOI:** 10.1101/2023.06.27.543376

**Authors:** Carlos Flores, Jefferson Ling, Amanda Loh, Ramón Garcia Maset, Angeline Aw, Ian J. White, Raymond Fernando, Jennifer L. Rohn

## Abstract

Urinary tract infection is among the most common infections worldwide, and is typically studied in animals and cell lines with limited uropathogenic strains. Here, we assessed diverse bacterial pathogens and asymptomatic bacteria (ASB) in a human urothelial microtissue model including full stratification/differentiation and urine tolerance. Several uropathogens and ASB-like *E. coli* invaded intracellularly, suggesting invasion is a shared survival strategy, instead of a virulence hallmark. The *E. coli* adhesin FimH was required for intracellular community formation, but not for invasion. Other shared lifestyles included filamentation (Gram-negatives), chaining (Gram-positives) and hijacking of exfoliating cells, while biofilm-like aggregates formed mainly with *Pseudomonas* and *Proteus*. Urothelial cells expelled invasive bacteria in Rab-/LC3-decorated structures, while highly cytotoxic/invasive uropathogens, but not ASB, disrupted host barrier function and strongly induced exfoliation and cytokine production. Overall, this work highlights diverse species-/strain-specific infection strategies and corresponding host responses in a human urothelial microenvironment, providing insights at the tissue, cell and molecular level.

**One-Sentence Summary:** A human urothelial model revealed shared colonization strategies between uropathogens and asymptomatic bacteria, and pathogen-specific innate immune responses

## INTRODUCTION

Urinary tract infection (UTI) is one of the most common human bacterial infections, with recurrence rates of ∼30% within 6 months highlighting the suboptimal performance of antibiotics*(1)*. Due to successive treatment rounds, including prophylaxis, UTIs exacerbate the global antimicrobial resistance crisis*(2, 3)*. Together with significant morbidity, reduced quality of life and mortality, UTI represents a hefty economic burden for healthcare systems worldwide*(1)*.

Uropathogenic *Escherichia coli* (UPEC) is the most common and well-studied uropathogen*(4)*, but while urothelial invasion and intracellular bacteria communities (IBCs) were defined in the murine model, relatively few papers have clearly observed intracellular bacteria in patient-derived specimens due to the difficulty inherent in such studies*(5)*. Hence, a key question in the field is how common intracellular invasion is in the human setting and how it is achieved.

Matters are even less certain for other UPEC besides UTI89 and CFT073, to say nothing of non-UPEC strains. The behaviour of asymptomatic bacteria (ASB) from the healthy urobiome, including *E. coli(6, 7)*, is also a major gap in the field. Moreover, other species are not rare, especially in older people, where polymicrobial infections abound, as well as in hospitals and catheterised patients, where *Pseudomonas aeruginosa*, *Proteus mirabilis, Enterococcus* sp. and *Klebsiella pneumoniae* thrive*(4)*.

Mouse models and bladder/kidney cancer cell lines have enabled advances in the understanding of host-uropathogen interactions, but key tissue ultrastructural and physiological species differences*(5)* have triggered the recent development of a few cell-based 3D urothelial models and organoids*(8–11)*.

Here, we deployed a robust 3D microtissue model recapitulating critical human urothelial features, such as urine tolerance, stratification and differentiation, to ask (1) whether UPEC, including clinical isolates and adhesin mutants, could invade this model and form IBCs; (2) how ASB-like *E. coli* colonize this environment; (3) whether the main Gram-positive and -negative uropathogens use conserved or distinct strategies; and (4) how human microtissue responds to both uropathogens and ASB. Our results revealed a surprising array of time-, species- and strain-specific diversity, with shared strategies between pathogens and ASB-like bacteria. Moreover, the host response usually correlated with specific bacterial strategies such as invasion or host barrier disruption. Overall, this model provides unique insights into human host-uropathogen interactions at both cellular and tissue levels.

## RESULTS

***(Higher resolution figures may be provided upon request)***

### Urothelial microenvironment promotes UPEC strain-specific morphology and invasion phenotypes

We generated 3D-UHU (<u>3D</u> <u>U</u>rine-tolerant <u>H</u>uman <u>U</u>rothelial) microtissue models (Fig. 1A,B), characterized in-depth in Jafari and Rohn, 2023*(11)*, recapitulating key human features (Fig.1C-E) including: i) thickness (∼60 μm) in stratification, *i.e.* 7-8 cell layers; ii) differentiation of the three main urothelium sublayers – basal, intermediate, and apical umbrella cells (expressing terminal differentiation markers – fig. S1A); iii) apical production of glycosaminoglycans (GAG layer); iv) robust barrier function; and v) 100% urine tolerance for long time periods. We first examined the behaviour of UP+EC during infection in urine over 24 h, testing well-known cystitis (UTI89) and pyelonephritis (CFT073) strains (Fig. 1F), alongside clinical UTI isolates (EC10, EC6, ClinA). All strains were mainly scattered on the apical urothelial surface 3 hours post-infection (hpi), while the population increased over time (data not shown). From 6 hpi, CFT073, UTI89 and EC10 began to invade (Fig. 2A), accompanied by the formation of large communities (IBCs) that could later erupt, more pronounced after 12 hpi (Fig. 2B), and with intracellular bacteria reaching >10^6^ CFU/mL (Fig. 2C). In contrast, ClinA and EC6 were able to invade but remained largely unaggregated (fig. S1C) as intracellular isolated bacteria (IIB) with lower loads of intracellular bacteria (≤ 10^6^ CFU/mL, Fig. 2C).

**Fig. 1.**
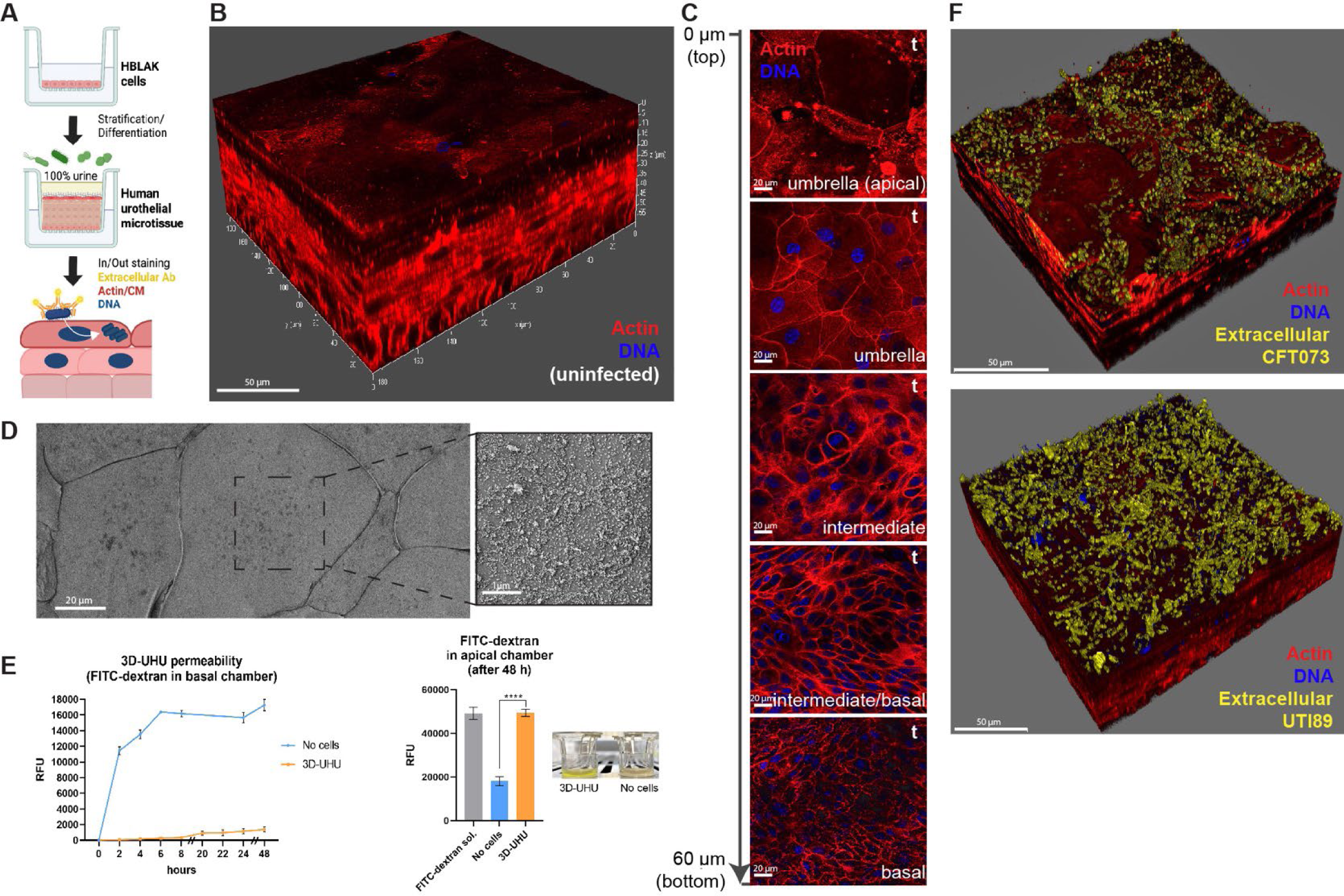
Establishment of the human urothelial infection model (3D-UHU). (**A**) Schematic representation of 3D-UHU development and staining for extra-*vs* intracellular bacteria. (**B**) 3D view of uninfected model. (**C**) 3D-UHU top-down (t) cross sections from top (0 μm) to bottom (60 μm) depicting the different cell morphology of the 3 sublayers (umbrella, intermediate, basal). Blue, DNA; Red, F-actin. (**D**) 3D-UHU topography showing tight junctions, GAG polysaccharides and microplicae/hinges on umbrella cells. (**E**) FITC-dextran (4 kDa) translocation assay using Tranwells inserts without cells and with 3D-UHU. Fluorescence was measured in the basal (left) and apical (right) chambers over 48 h (****, *p* < 0.0001), the latter was compared with FITC-dextran stock solution. Image showing FITC-dextran retention by 3D-UHU insert after 48 h (in bright yellow). (**F**) 3D view of 3D-UHU 12 hpi with UPEC CFT073 and UTI89 (in yellow). Confocal (B,C,F) and SEM (D) images representative of a minimum of 4 independent biological replicates per strain.

**Fig. 2.**
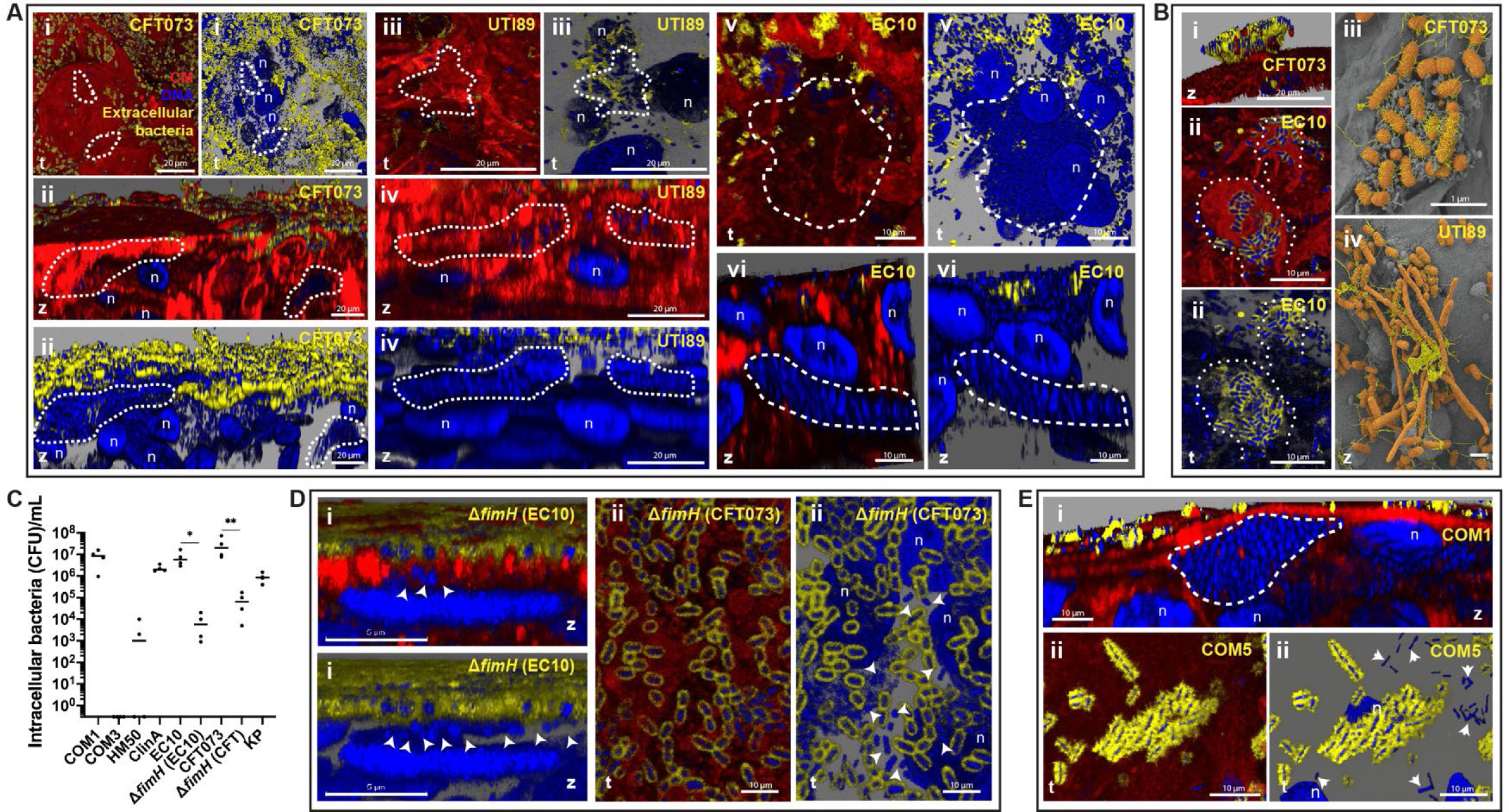
Invasion and IBC formation by UPEC and ASB-like *E. coli.* (**A**) Invasion and IBC formation (dashed lines) by UPEC CFT073 (i,ii), UTI89 (iii,iv) and EC10 (v,vi). t, top-down view; z, side view. Yellow, extracellular bacteria; Blue, DNA of host nuclei (n) and bacteria (intra and extracellular); Red, cell membrane (CM). (**B**) UPEC IBC eruption at 12 hpi, with sprouting filaments in UTI89 (iv). (**C**) Intracellular bacteria quantified by gentamicin protection assay 12 hpi with ASB (COM1, 3 and HM50), UPEC (ClinA, EC10, CFT073 and respective Δ*fimH* mutants), and *K. pneumoniae* (KP) at MOI 100 (inoculum of 10^6^ bacterial cells). Data plotted as mean of 4 independent biological replicates, **, *p* < 0.01; *, *p* < 0.1. (**D**) Invasion without IBC formation by Δ*fimH* (arrowheads), in EC10 (i) and CFT073 (ii) backgrounds. (**E**) Invasion by ASB COM1 with IBC formation (i, dashed line), and COM5 as IIB (ii, arrowheads), at 12 hpi. Confocal (A,Bi-ii,D,E) and SEM (Biii-iv) images representative of a minimum of 4 independent biological replicates per strain.

Urothelial damage caused by UPEC matched their invasive behaviours, with CFT073, closely followed by UTI89, having the strongest impact: a ∼6-fold increase in barrier permeability (Fig. 3A), and ∼20% more cytotoxicity (Fig. 4A). In contrast, clinical UPEC isolates EC6 and ClinA had a weaker impact on both cytotoxicity (∼8%) and permeability.

**Fig. 3.**
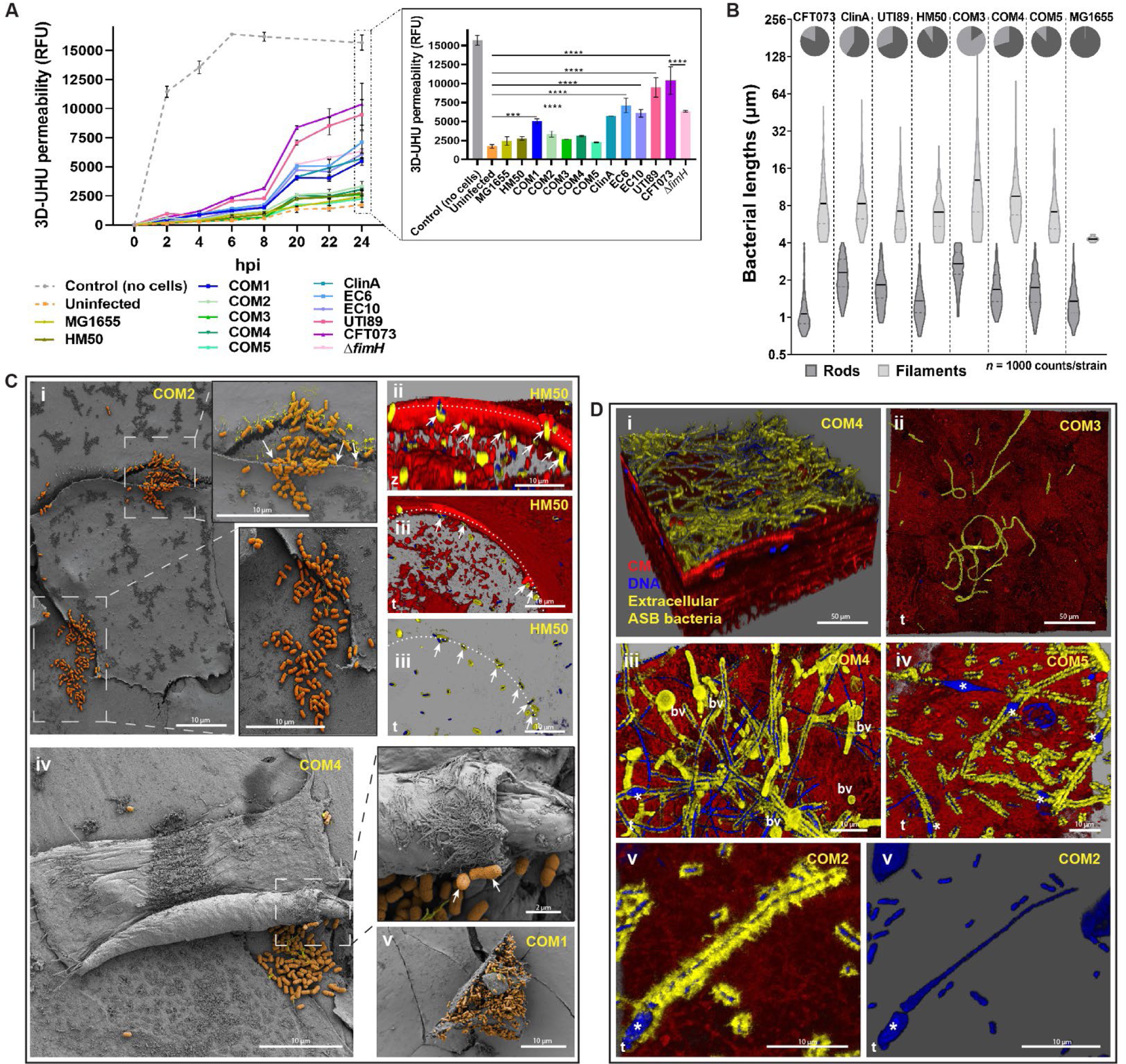
UPEC and ASB-like *E. coli* colonization strategies and morphology in the 3D-UHU. **(A)** 3D-UHU barrier function assessed by FITC-dextran (4 kDa) permeability assay before/after infection. Fluorescence measured in basal chambers over 24 h; data plotted as mean ± SE of biological triplicates. Inset compares the final time point, ****, *p* < 0.0001; ***, *p* < 0.001. (**B**) Size distribution of *E. coli* after 12 hpi. Black lines - means of rod (dark grey) and filament (light grey) sizes; Pie charts - proportion of each group in *n* = 1000 bacteria per strain. (**C**) Adhesion by ASB COM2 (i), HM50 (ii-iii) and COM4 (iv) to underside of exfoliating cells. Dotted lines (ii-iii) depict edge of cell with bacteria underneath (arrows). Adhesion by COM1 (v) to a dying cell. t, top-down view; z, side view. Staining as in Fig. 2. (**D**) Filamentation by ASB (i-v) accompanied by bacterial outer membrane vesiculation (iii, Bv) and fusiform nuclei (iii-v, *) at 12 hpi. Confocal (Cii-iii, D) and SEM (Ci,iv,v) images representative of a minimum of 4 independent biological replicates per strain.

**Fig. 4.**
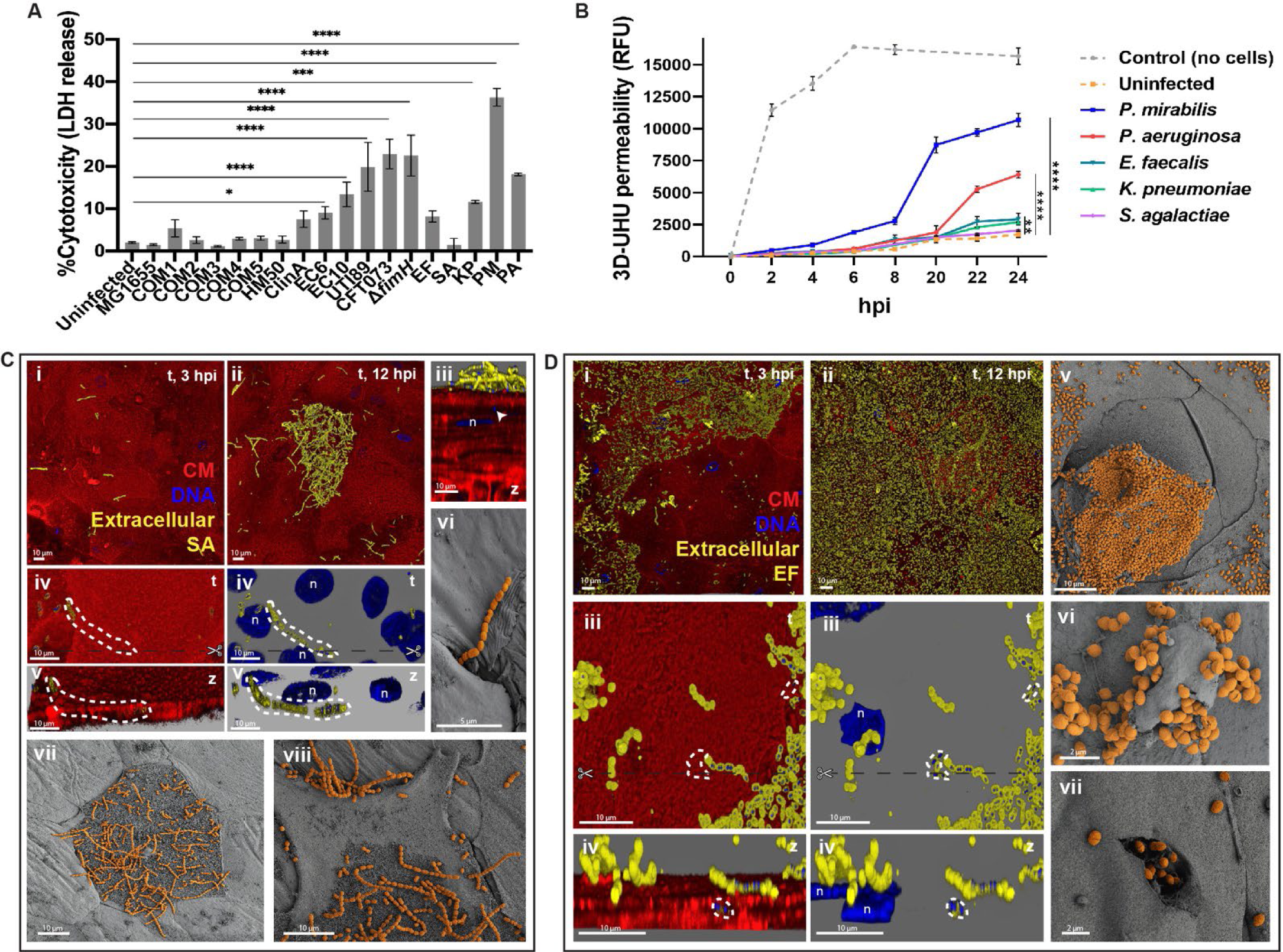
Infection strategies by non-UPEC uropathogens in the 3D-UHU. (**A**) Cytotoxicity caused by uropathogens and ASB 12 hpi, assessed by LDH release assay. Data plotted as mean ± SE of biological triplicates, ****, *p* < 0.0001; ***, *p* < 0.001; *, *p* < 0.1. (**B**) 3D-UHU barrier function post-infection with non-UPEC uropathogens, assessed by FITC-dextran (4 kDa) permeability assay. Fluorescence measured in basal chambers over 24 h. Data plotted as mean ± SE of biological triplicates (****, *p* < 0.0001; **, *p* < 0.01). (**C**) *S. agalactiae* (SA) in discrete regions of the urothelial surface (i,ii,vii,viii), associated with damaged upper cell host membranes (vii,viii). Cocci invasion (iii, arrowhead), and chains underneath upper cell layers (iv-vi). Scissors (iv) depict the place of cross section for the side view. (**D**) *E. faecalis* (EF) spread on the urothelial surface (i,ii), and chains translocating between the upper cell layers (iii-iv, dashed lines). Scissors (iii) depict the place of cross section for the side view. Heavily colonized umbrella cells being exfoliated (v) and cocci erupting from a vesicle-like structure (vi) and urothelial cell (vii). t, top-down view; z, side view. Staining as Fig. 2. Confocal (Ci-v, Di-iv) and SEM (Cvi-viii, Dv-vii) images representative of a minimum of 4 independent biological replicates per strain.

Elongated cells and filaments were also found for all UPEC (Fig. 3B), sometimes accompanied by the eruption of IBCs (Fig. 2Biv). Small biofilm-like surface clusters could sporadically be observed (fig. S1B).

### UPEC FimH is important for IBC formation but not essential for invasion

Despite being severely impaired, significant invasion still occurred in UPEC knockouts of the FimH adhesin (∼10^4^ CFU/mL) with distinct O-antigen serotypes CFT073 (O6) and EC10 (O25b) (Fig. 2C), but neither formed large IBC (Fig. 2D). Moreover, filamentation was not impaired in any mutant, but CFT073 Δ*fimH* had significantly enhanced biofilm production, the highest among pathogenic and ASB *E. coli* tested (fig. S2A). This mutant had also an impaired effect on 3D-UHU permeability (∼3 to 4-fold increase, Fig. 3A), but caused similar cytotoxicity compared with the parental strain (Fig. 4A).

### Asymptomatic bacteriuria (ASB)-like *E. coli* share survival strategies with UPEC, including invasion, filamentation and hijacking exfoliating cells

Intracellular invasion of UPEC has long been considered a hallmark of virulence in mice and cell lines*(12)*. We interrogated this in our human microtissue model with lab strain MG1655, the well-characterised ASB strain HM50 and other 5 genetically distinct ASB-like *E. coli*: COM1-5, recovered from urine of healthy volunteers. COM1 and COM5 could invade umbrella cells, albeit differently: COM1 invaded frequently (∼10^7^ CFU/mL, Fig. 2C) with IBC formation comparable to some UPEC (Fig. 2Ei), while COM5 invaded only sporadically, and remained isolated (Fig. 2Eii and fig. S1D). We also detected very rare invasion events by HM50, mostly in exfoliating cells, ranging from 0 to ∼10^3^ CFU/mL (Fig. 2C). This finding also supports FimH not being essential for invasion, as HM50 does not produce a functional type 1 pili*(13)*. Most of the other ASB-like bacteria were also sparsely piliated (fig. S3). COM2-4 and MG1655 were only observed on the apical surface (e.g. in Fig. 3C,D), with no intracellular COM3 detected by gentamicin protection assay (Fig. 2C).

On the other hand, filamentation was extremely common among ASB-like strains, observed from 3 to 24 hpi, even in non-invasive strains (Fig. 3B,D). Mean bacterial lengths (∼1-2 μm rods and ∼8-10 μm filaments) were similar among UPEC and ASB, but with strain-specific variation in the proportion of filaments *vs* rods (Fig. 3B). For COM3, the bacterial population was almost entirely filamentous (∼85%), with longer mean sizes (> 3 μm rods and > 9 μm filaments), but they poorly adhered and were easily washed away. Despite encoding the FimH adhesin, COM3 probably lacks a functional type 1 pili and other fimbriae due to mutations detected in *fim* and other operons (data not shown), similarly to HM50, and as supported by their lack of hemagglutination ability (fig. S4). Moreover, ASB filamentation was often accompanied by nucleoid enlargement, e.g. into a fusiform shape, (Fig. 3Diii-v) and outer membrane vesiculation (Fig. 3Diii).

Unlike UPEC, most ASB-like *E. coli* caused negligible damage to 3D-UHU, with the exception of the IBC-forming COM1, inducing moderate loss of barrier permeability (Fig. 3A), but no significant cytotoxicity (Fig. 4A). However, a common phenotype among most *E. coli* was adherence/colonization of the underside of cells being exfoliated and/or dying (Fig. 3C).

### Urothelial microenvironment elicits species-specific behaviours in non-UPEC uropathogens

Given the unexpected diversity seen within *E. coli*, we next inspected the behaviour of other common uropathogens: *E. faecalis* (EF), *S. agalactiae* (SA), *K. pneumoniae* (KP), *P. aeruginosa* (PA), and *P. mirabilis* (PM). First, we detected that, even more dramatically than UPEC, PM impaired urothelial barrier function by ∼84% alongside a ∼35% increase in cytotoxicity (Fig. 4A,B). PA and KP caused less cytotoxicity (18 and 14%), but the response was comparable to most UPEC, followed by EF, while SA showed the weakest impact on 3D-UHU (Fig. 4A,B). In addition, all pathogens colonised vast areas of the urothelial surface, except SA (Fig. 4C) and KP (Fig. 5A), which were sparsely distributed even after long periods. KP was found in host membrane ruffles or surrounded by spike-like protrusions (Fig. 5Aiii,iv,vi-viii). With SA, discrete chain niches were found after 6 hpi, as small aggregates or on top of umbrella cells with damaged plasma membranes (Fig. 4C and fig. S2C). In fact, the formation of chains was a common strategy among uropathogens, although with time- and species-dependent heterogeneity. Gram-positives formed longer chains from 3-12 hpi with increasing amounts of isolated cocci over time (Fig. 4C and D, i,ii), while PA (Fig. 5Bi,ii) and PM chains (Fig. 5Ci,ii) tended to be longer after 12 hpi, possibly supporting the development/connection of biofilms, as previously reported in different infection contexts*(14, 15)*. In addition, both Gram-positive (Fig. 4Civ-vi, Diii-iv) and PM (Fig. 5Ciii) chains were able to translocate paracellularly, mainly between cells undergoing exfoliation, and in case of PM, access deeper layers and reside in between/inside intermediate cells (Fig. 5Ciii). For PA, long chain formation coincided with the development of larger biofilm-like aggregates that merged and thickened, often containing cell debris and covering wide portions of the urothelial surface by 12 hpi (Fig. 5Bii,vi-viii and fig. S2E). This was accompanied by the decrease of PA size, from ∼ 4 down to ∼1 μm (Fig. 5Biii). In addition, PA and PM biofilms usually contained precipitates with a crystalline aspect, and in the case of PM, these deposits were particularly larger, more abundant and interjunctionally positioned (Fig. 5Cvi,viii and fig. S2D). Moreover, PA and PM produced more biofilms in urine or optimal media alone compared with the other uropathogens, approximately >6x higher for PA, despite the general negative impact of urine on biofilm production (fig. S2A).

**Fig. 5.**
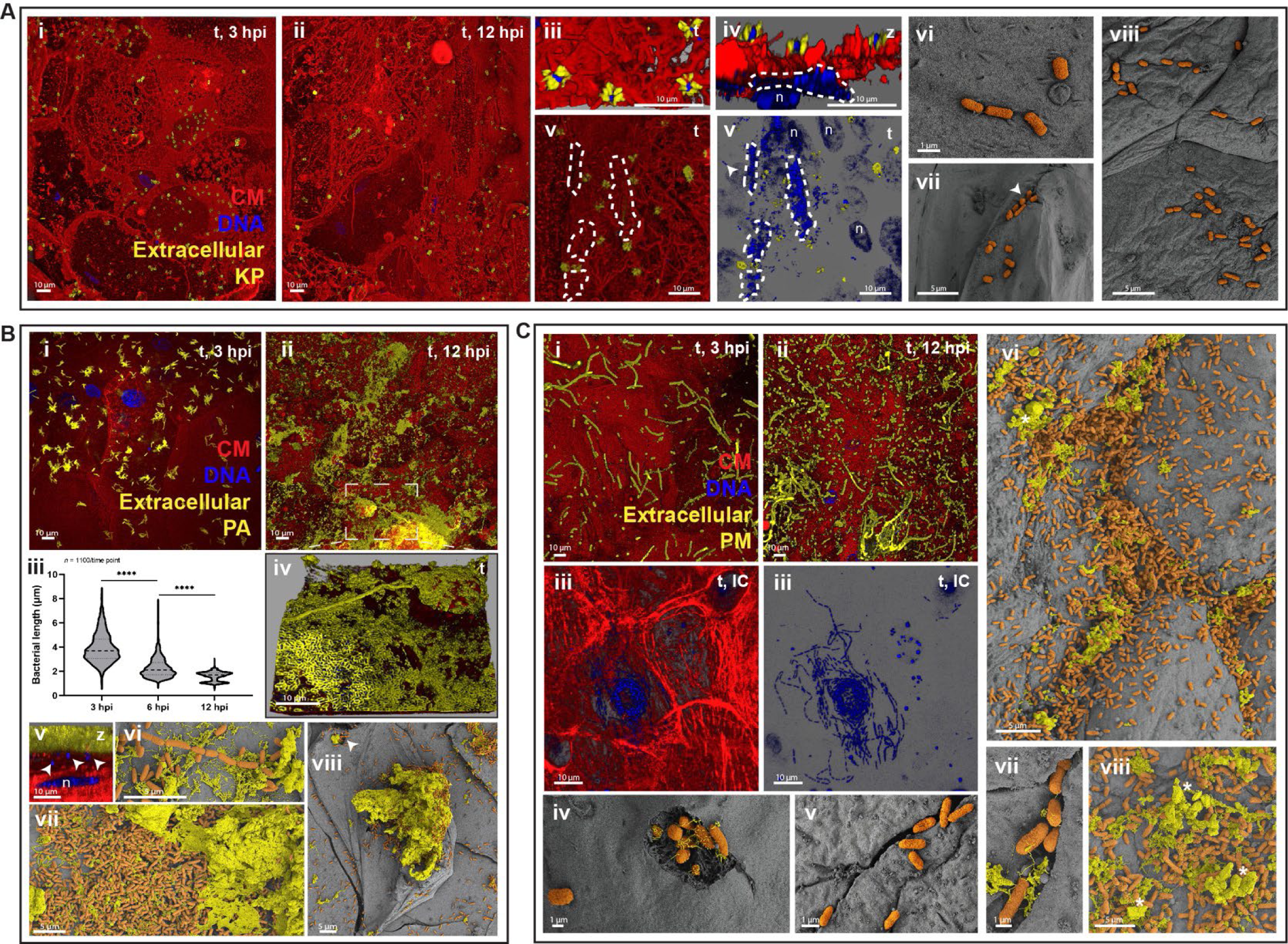
Infection strategies and morphology of Gram-negative non-UPEC uropathogens in the 3D-UHU. (**A**) *K. pneumoniae* (KP) infection. Bacteria scattered on the urothelial surface (i-ii); membrane ruffling and spike-like structures surrounding KP (iii-viii). Dashed lines, IBC (iv,v); Arrowhead, possible eruption (vii). t, top-down view; z, side view. Staining as in Fig. 2. (**B**) *P. aeruginosa* (PA) infection. Formation of biofilm-like aggregates (i,ii,iv,vi-viii), incorporating cell debris and precipitates (vi-viii). Violin plot showing decrease in bacteria length over course of infection (iii, *n* = 1100 bacteria/time point; ****, *p* < 0.0001). Arrowheads, intracellular bacteria (v,viii). (**C**) *P. mirabilis* (PM) infection. Rods, chains and/or elongated forms on 3D-UHU surface (i-iii), inside a cross-section of intermediate cell layers (IC, iii), inside umbrella cells (iv), and penetrating paracellularly in inflamed urothelium (v-vii). Interjunctional biofilms with crystalline precipitates (*,vi-viii). Confocal (Ai-v, Bi-v, Ci-iii) and SEM (Avi-viii, Bvi-viii, Civ-viii) images representative of a minimum of 4 independent biological replicates per strain.

Intracellular invasion was commonly observed for KP and PM, as for UPEC after 6 hpi, including in intermediate cells for PM after paracellular penetration (Fig. 5Ciii-vii), and with formation of large IBCs in umbrella cells by KP (Fig. 5Aiv-v). The remaining uropathogens could also invade, but it was rare and only seen after 12 hpi (e.g. Fig. 4Ciii, Dvi-vii, 5Bv).

Similar to *E. coli*, adhesion to the underside of cells undergoing exfoliation/dying was common among all non-UPEC uropathogens (Fig. 4Dv, 5Bviii and fig. S2Bii, Civ-v).

### A battery of tissue innate and autonomous urothelial responses differentially target invasive *vs* extracellular bacteria

We frequently observed the extrusion of host vesicle-like structures and/or cell fragments of varying sizes from umbrella cells during infections (Fig. 6A and fig. S5A). With SA, these structures were usually devoid of bacteria, but heavily surrounded by long chains (fig. S5A). In contrast, infections with more invasive strains (e.g. CFT073, UTI89 and COM1) featured bacteria inside them, often localized in areas with extensive plasma membrane ruffling (Fig. 6A and fig. S5B,C).

**Fig. 6.**
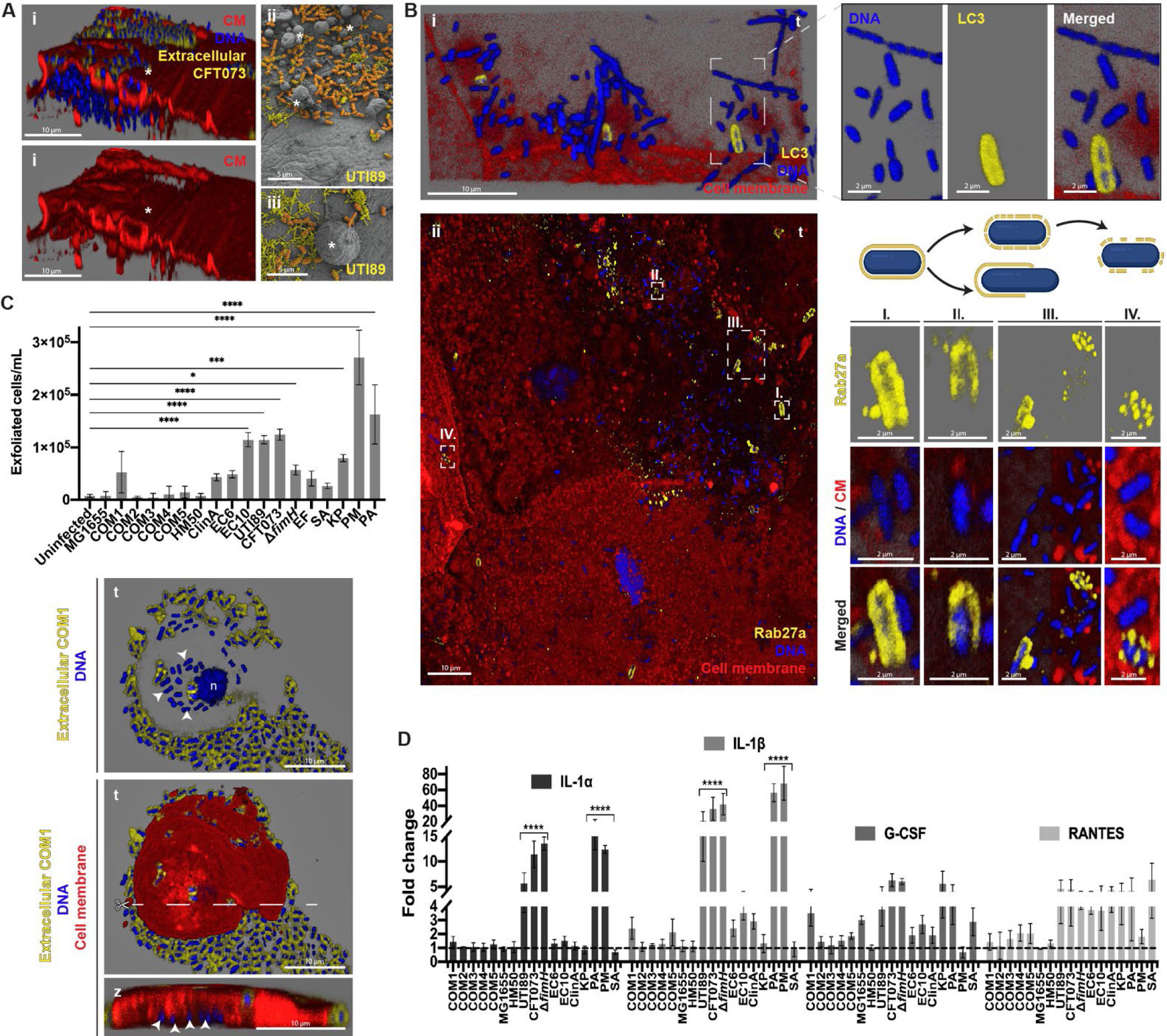
Host responses to uropathogens and ASB-like bacteria. (**A**) Membrane ruffling and formation of blebs/vesicle-like structures (*) by umbrella cells 12 hpi with CFT073 (i) and UTI89 (ii,iii). (**B**) Expelled CFT073 in exosome-like structures decorated with LC3 (i) and Rab27a (ii) (yellow). I-IV depict different encasement patterns suggesting bacterial release either through elongation or case degradation (scheme). Staining as in Fig. 2. t, top-down view; z, side view. (**C**) Host cell exfoliation 12 hpi with uropathogens and ASB, and exfoliated cell with intracellular COM1 (arrowheads). Scissors depict cross section placement for the side view. Data plotted as mean ± SE of biological triplicates, ****, *p* < 0.0001; ***, *p* < 0.001; *, *p* < 0.1. (**D**) Cytokine (IL-1α, IL-1β, G-CSF and RANTES) production by 3D-UHU 12 hpi. Fold changes compared to uninfected controls (dashed line) plotted as mean ± SE of biological triplicates, **** (*p* < 0.0001). Confocal (Ai,B,C) and SEM (Aii,iii) images representative of a minimum of 4 independent biological replicates per strain.

We also detected encasement of UPEC in “exosome-like” structures containing the phagosome marker LC3 (Fig. 6Bi). In addition, Rab27a (related to exocytosis and vesicle trafficking) also colocalized with some extracellular UPEC, ASB-like *E. coli* and KP (Fig. 6Bii). These structures were not homogeneous, varying from a complete “cage” to porous structures or protein patches (Fig. 6Bii). Rab27b (a marker of fusiform vesicles) was also observed in these structures, but as it was highly expressed throughout the umbrella cell layer, specificity for encased bacteria was not so obvious (data not shown).

Sporadic umbrella cell exfoliation, a homeostatic urothelial mechanism*(16)*, could be reproduced by uninfected 3D-UHU (Fig. 6C). However, more exfoliation occurred in infection with aggressive strains, such as with the invasive PM (∼3x10^5^ exfoliated cells/mL), CFT073, UTI89 and EC10 (∼1x10^5^ cells/mL); and the biofilm former PA (∼1.5x10^5^ cells/mL). With IBC formers, these exfoliated cells were frequently full of bacteria (Fig. 6C). In contrast, excess exfoliation did not occur for ASB-like strains, even the efficient invader COM1.

The apparent specificity in host responses against uropathogens was also extended to cyto/chemokine production (Fig. 6D). Among the 16 analytes assessed, IL-1β production was by far the most accentuated, especially with UTI89, CFT073, PA and PM (>10-fold higher), followed by IL-1α, G-CSF and RANTES. Increase in IL-6, IL-8, and Pentraxin 3 against uropathogens was also detected (fig. S5D). The pro-inflammatory reaction agrees with the distended aspect of upper cell layers suggestive of urothelial inflammation, e.g. in PM (Fig. 5Cvi and fig. S2Diii). Surprisingly, infections with PM or PA also induced similar or much lower levels of some cytokines compared with ASB or uninfected controls, with IL-6 production being particularly affected (fig. S5D). Moreover, in agreement with the abovementioned data, CFT073 and UTI89 triggered the most pronounced response among UPEC, while most ASB-like *E. coli* induced a response similar to uninfected controls, the weakest being HM50 and COM3 (Fig. 6D and fig. S5D). No significant changes were observed for Lactoferrin and Lipocalin-2, whereas osteopontin and IFN-γ were not detected.

## DISCUSSION

Two decades after the first report of intracellular UPEC in the murine UTI model*(17)*, many questions remain about host-pathogen interactions in the human bladder. Driven by the need for advanced human-derived models*(5)*, we developed 3D-UHU as a robust model mimicking critical human urothelial features. The long-term tolerance of urine is key, as it affects not only host cells*(18)*, but also growth, metabolism*(19)*, pathogenicity, e.g. biofilm formation (Fig. 2SA; *(14)*) and virulence factor expression*(20)*. Moreover, the multiple intermediate cell layers, as opposed to just one in mice*(21)*, are critical for assessing the establishment of deeper bacterial reservoirs and regeneration after exfoliation, as well as the balance between recurrence and UTI resolution in humans.

Using this human-relevant microenvironment, with undiluted urine, we tested various uropathogens and ASB-like bacteria, revealing a surprising diversity of time-, species- and strain-specific behaviours compared with what is documented for animal and *in vitro* models, which typically featured a more restricted set of UPEC strains. In mice, UPEC cause stereotypical IBCs*(17)*, which could be recapitulated in 3D-UHU with some clinical isolates. However, two other clinical strains invaded without forming IBCs, remaining as IIB. In addition, the ASB-like strain COM1 invaded frequently and formed large IBCs, while other ASB-like also invaded, albeit rarely. These results suggest that invasion represents a shared strategy for persisting in a harsh environment, not necessarily a virulence trait.

Adherence and invasion in the absence of functional FimH, a major player during mouse UTI, suggests some redundancy with other virulence factors, which is supported by the increasing list of such prominent factors and the tight coordination of their expression*(22, 23)*. Furthermore, these data highlight the importance of developing alternative drugs that target multiple virulence factors to supplement promising anti-FimH therapeutics*(24)*. In contrast, no IBC formation was observed in FimH’s absence, stressing its importance in intracellular bacteria-bacteria interactions, and agreeing with the crucial role of type 1 pili in both IBC initiation and maturation in mouse bladders*(25)*.

Invasion has also been reported for most common non-UPEC uropathogens in mice or cell lines*(18, 26–28)*. In 3D-UHU, invasion was prominent with KP, which formed IBCs similar to those of UPEC, and present but rare in other species. In addition, host membrane spike-like protrusions surrounded individual KP on the surface, resembling internalization pathways described for *Salmonella* or *Shigella(29)* that confer high invasion efficiency even with weaker adhesion/colonization (Fig. 5 and Rosen et al.*(26, 30)*).

The efficient way that PM infiltrated both paracellularly and intracellularly supports previous findings in cell lines and mice, for bladder and kidney*(31–33)*. We showed that this can occur in deeper intermediate cells, which is difficult to show in thinner mice urothelia. However, as previously suggested*(33, 34)*, invasion was not the dominant PM lifestyle, nor for PA, where extensive biofilm clusters were formed with mineral deposition and/or cell debris. These aggregates resembled initial stages of encrustations and bladder stones, commonly detected in catheter-associated UTI animal models resulting from ureolytic biomineralization*(33, 35)*. Our data suggest that these crystalline biofilms can occur even in the absence of indwelling devices or neutrophil recruitment/extracellular traps.

On the other hand, some UPEC, SA and PA could not so efficiently invade the urothelium (and/or replicate/persist). In these cases, IIB might be trapped in intracellular compartments, easier targets for urothelial expulsion via exocytosis in exosome-like structures*(36, 37)*. In addition to the previously reported LC3 and Rab27b decorating these “cages”, we showed that Rab27a can also colocalize with expulsed bacteria. Rab27a is still not widely studied in the infection context, with moderate to absent expression reported in uninfected urothelium from different species*(38–40)*. Here, its distinct labelling pattern (from uniform to porous) suggests a bacterial escape mechanism either via cell elongation, or cage degradation, as for a recent study showing that intracellular UPEC sense engulfment in fusiform vesicles, triggering expression of enzymes to escape and form IBCs*(41)*.

When more subtle host defences fail, the expulsion of bacteria in host cell fragments or wholesale cell exfoliation can serve as “last resort” measure for highly invasive*(42)* and/or cytotoxic UPEC strains*(43)*. This agrees with the particularly marked response we observed for CFT073, but also against PM and PA, accompanied by barrier disruption and extensive cell membrane ruffling. However, this strategy may backfire for the host, as it allows establishment of deeper reservoirs, paracellular translocation and/or cell death (as commonly observed here with PM); hence, bacteria may boost this response deliberately *(32, 43–45)*. We observed that most bacteria took advantage of the exfoliation process to adhere underneath cells, inserting between the upper layers. For PA, this strategy contributed to the incorporation of cell debris, possibly to thicken their protective biofilm aggregates, which was previously shown, not only for PA, in distinct infection contexts*(46, 47)*. Moreover, the “highjacking” behaviour may provide temporary shelter from luminal defences, while exploiting the sudden exposure of host adhesive molecules or matrix components. This may help to explain how poorly adherent and avirulent ASB, e.g. HM50, can persist and thrive in the urothelium despite flow*(48, 49)*. The formation of biofilms, associated with lack of host defence activation, has been the key hypothesis for ASB persistence and was also observed here*(13, 49)*. In addition, our data support the recently reported “rolling-shedding” colonization mechanism, in which UPEC hijacks exfoliated cells to spread and persist throughout the urothelial surface*(50)*. Moreover, host cells in poor physiological status are easier targets for toxins and siderophores, providing a useful source of nutrients, otherwise limited in urine*(4)*.

Chaining was also extremely common for most bacteria, apart from UPEC, KP and ASB-like *E. coli*, where cell elongation and filamentation predominated. Although filaments have been studied in the context of UPEC IBC eruption and immunity evasion*(51, 52)*, our data suggest they are also important for survival in the harsh human bladder microenvironment, and not solely associated with pathogenicity, in agreement with recent evidence*(53)*. These bacteria also showed other alterations previously associated with growth in stressful environments, such as outer membrane vesiculation or cell/nucleoid enlargement*(54, 55)*.

In our model, more aggressive strains induced the production of several proinflammatory cytokines, reminiscent of observations in mouse bladders and UTI patients*(56)*. The opposite was observed for ASB, with the poorly adherent COM3 and HM50 eliciting the lowest levels. The particularly enhanced levels of IL-1β against highly cytotoxic strains agrees with previous studies where UPEC activate the host inflammasome in a haemolysin-dependent manner*(57–59)*. This enhancement was not necessarily related to invasion efficiency, as PA and CFT073 Δ*fimH* induced higher levels compared with more invasive strains. Other toxins most likely contribute to the response to PM and PA. For instance, in non-UTI contexts, PA pore-forming toxins also induced inflammasome pathways*(60)*. In addition, cumulative effects of IL-1β inflammasome-independent production pathways, not addressed here, might also occur*(57)*. In contrast, the non-hemolytic SA used here induced lower levels of IL-1β compared with haemolytic-positive strains*(58, 61)*.

The downregulation of major cytokines after PM or PA infection (namely IL-6) was notable, considering the extensive damage and cytotoxicity caused. Mechanisms associated with timing of infection or specificity in host immunity might underly these particular profiles. In addition, bacterially active immunomodulation (e.g. via outer membrane vesicles) is also possible*(62)*, but still largely unexplored in the UTI context.

Altogether, the 3D-UHU microtissue model, as a good facsimile of the human urothelial microenvironment, revealed a wide variety of bacterial lifestyles and host responses, which highlights the need to develop novel therapeutics counteracting the current “one-size-fits-all” UTI treatment approach. Strategies such as filamentation, invasion and host hijacking may aid persistence of both uropathogens and ASB, but are not necessarily always related with virulence, reinforcing the need for more studies of the urobiome*(7)*.

Several avenues still remain to be (re)addressed in human models in order to be used in clinics (e.g. vascularization), but 3D-UHU should prove a tractable and robust complement to animal studies that can be fine-tuned (e.g. with biomechanical stimuli or immune cell co-culture) to better understand human UTI, or even used as a powerful platform for drug development, while providing details at tissue, cellular and molecular levels.

## MATERIALS AND METHODS

### Study Design

The initial goal of this study was to characterize critical events during uropathogenesis and host responses in a human-like urothelial microenvironment, with diverse UPEC clinical isolates and asymptomatic bacteria. We hypothesized that invasion and intracellular communities could occur in different ways depending on the bacteria. We also wanted to know if FimH, the main UPEC adhesin used as potential therapeutic target, demonstrates a critical role in a human tissue-like scenario. Using our new human microtissue urothelial model comprising barrier function, stratification/differentiation and urine tolerance for long periods, we infected the bacteria during different periods 0-48 h and at distinct MOI (10-100). Bacterial expression of the type 1 fimbriae was verified by hemagglutination assays. At fixed time points, we observed the progression of infection/colonization using confocal and SEM. Images were representative of a minimum of 4 independent biological replicates per strain. We also performed antibiotic protection assays to determine bacterial invasion efficiency intracellularly. We investigated bacterial morphological alterations and quantified the rods-to-filaments ratio after infection in a blind automated manner. To assess the effect of the bacterial strategies in the host, we determine the induced cytotoxicity by LDH release from the microtissue, and its barrier permeability over time using a FITC-dextran (4 kDa) diffusion assay. Using microscopy, we addressed the expulsion of bacteria in exosome-like structures, host vesiculation/blebbing and exfoliation, while the apical milieu was used to quantify cyto/chemokines production and exfoliated cells. Ethical approval for collection and use of isolates from UTI patients and healthy volunteers were in place. Given the diversity of urothelium-*E. coli* interaction, we used a similar approach to assess infection strategies by other common Gram-positive/-negative uropathogens and respective host responses.

### Human cell line

Spontaneously immortalised, non-transformed human bladder epithelial cells (HBLAK, CELLnTEC) were supplied containing approximately 5×10^5^ cells per vial and were kept in liquid nitrogen until further use. They were maintained in prewarmed CnT-Prime medium (CELLnTEC), in humidified environment at 37°C and 5% CO_2_ prior to passaging. Passages were performed using Accutase^TM^ (Sigma-Aldrich) for detachment of 80-90% confluent cells, incubated for 7 minutes at 37 °C, after washing with Ca^2+^ and Mg^2+^-free phosphate buffered saline (PBS, GIBCO).

### Bacterial strains

The *E. coli* strains 83972 (HM50, asymptomatic bacteria) and CFT073, *P. aeruginosa* PAO1 (BAA-47), *S. agalactiae* G19 (13813) and *P. mirabilis* 7570 (51286) were obtained from the American Type Culture Collection (ATCC), while *K. pneumoniae* TOP52 and UPEC UTI89, recovered from patients with acute cystitis, were kindly provided by Scott Hultgren (Washington University St. Louis, USA), and *E. coli* MG1655 by Dr. Meriem El Karoui (University of Edinburgh). The isolates EC6 and EC10 are MLST ST131 serotype O25b bladder infection isolates*(63)* from Pfizer (Dept. of Vaccine Design, Immunology & Anti-Infectives Pearl River, NY, US). Gene knockout mutants of *fimH* were generated from CFT073 and EC10 parental strains using the λ-Red mediated homologous recombination system as previously described*(64)*. Clinical isolates from Royal Free Hospital, UK were *E. faecalis* (EF36), a previously reported clinical isolate from a patient with chronic UTI*(18)*; UPEC ClinA, isolated from a chronic UTI patient^57^, and five “ASB-like” *E. coli* isolates (COM1-5) were recovered from clean-catch midstream urine samples of healthy individuals and reported previously*(65)*. All strains were kept in glycerol at -80 °C until further use. They were maintained at 37 °C in optimal media before experiments: Luria broth for *E. coli* (LB, Sigma-Aldrich), tryptone soya broth (TSB, Oxoid) for *P. aeruginosa* and *S. agalactiae*, *P. mirabilis* and *E. faecalis*, and nutrient broth (NB, Oxoid) for *K. pneumoniae*.

### Generation of 3D human urothelial model (3D-UHU)

3D-UHU were generated as previously described*(11)*. Briefly, between passages 8 to 12, 80-90% confluent cells were detached as above described and 3 x 10^5^ cells/mL in pre-warmed CnT-Prime were seeded onto 12 mm, 0.4 μm pore polycarbonate filter (PCF) membranes in plastic inserts standing in 12-well Transwell plates (Corning^TM^), while the basal chamber was filled with 1.5 mL of the same media. After 2 days of incubation at 37 °C, 5% CO_2_, media in both apical and basal chambers was replaced by calcium-rich (1.2 mM) differentiation barrier medium (CnT-Prime-3D medium, CELLnTEC), designated day 0. After overnight incubation, media in the apical chambers was replaced with commercially available filter-sterilised human urine from 10 individuals (both sexes) per pool (BioIVT), while fresh media was replaced in the basal chamber. Urine/3D media changes every 3 days were performed until day 18 to 20, when models were used for subsequent experiments.

### Bacterial inoculations

Prior to infection, *E. coli* strains were grown for two consecutive overnight incubations statically at 37 °C, while one overnight incubation was used for the other species of bacteria. Bacterial numbers were quantified using QUANTOM Tx™ Microbial Cell Counter (Logos), according to the manufacturer’s instructions. For infection, bacteria were inoculated in the apical chamber using multiplicity of infection (MOI) 10 to 100 in urine, depending on the experiment. For reproducibility purposes, MOI was calculated on the basis of approximately 30,000 umbrella cells on the mature microtissue apical surface, as estimated by surface area and cell size. CnT-Prime-3D media was replaced in the basal chamber.

### Hemagglutination assays

Bacteria were grown in LB broth statically at 37 °C for two consecutive overnight incubations, and then diluted to OD_600nm_=0.5. Cells were harvested by centrifugation at 4000 x*g* for 5 min and resuspended in PBS or in 1% mannose (to inhibit type I fimbriae binding). Suspensions were then mixed with 5% or 1% (v/v) of guinea pig erythrocytes. Hemagglutination was visually monitored over 0-4h of incubation at 4 °C in microtiter wells, or imaged in glass slides after 30 min incubation at room temperature, using a Leica inverted microscope DMi1 (Leica Microsystems).

### Biofilm quantification assay

After growth as aforementioned, the formation of biofilms from all bacterial cultures was assessed in LB and 25% urine. Bacterial suspensions were adjusted to mimic MOI 15 and 30, onto Calgary biofilm devices (Innovotech), which are microtiter 96-well plates with lids that have pegs extending into each well. After static incubation at 37 °C, 5% CO_2_ for 24 h, the lids were removed and air dried for 20 minutes. The pegs were then submerged into a new plate containing 200 μL of crystal violet stain and incubated at room temperature for 30 min. Subsequently, the pegs were gently rinsed with distilled water and air dried before being submerged in 200 μL of 33% acetic acid (Sigma) for 15 min. Absorbance of the dissolved biofilms was measured using a Tecan microplate reader at OD_550nm_.

### FITC-dextran barrier permeability assay

To monitor the 3D-UHU barrier integrity upon infection, bacteria were inoculated at MOI 15 in 1 mg/mL of fluorescein isothiocyanate–dextran 4 kDa (FITC-dextran, Sigma-Aldrich) dissolved in CnT-Prime-3D. An empty Transwell and an uninfected model served as controls. After 0, 2, 4, 6, 8, 20, 22 and 24 h post infection (hpi), 50 µL of basal medium was transferred into a 96-well clear-bottom black polystyrene microplate (Corning^TM^). Fluorescence intensity was measured using a fluorescence plate reader (Tecan) at 490 nm excitation and 520 nm emission values.

### LDH cytotoxicity assay

Cytotoxicity caused by the bacteria at MOI100 after 12 hpi in 3D-UHU was assessed by quantification of the amount of lactate dehydrogenase (LDH) released into the apical milieu, using the commercially available CyQUANT™ LDH Cytotoxicity Assay (Thermo Fisher) as recommended by the manufacturer.

### Gentamicin protection assay

After 12 h post infection at MOI 100 (initial inoculum of 10^6 bacterial cells), the media in the apical chambers was replaced by 500 μg/ml gentamicin (in 100% urine), 150-250x above MIC (performed as described in Wiegand et al.*(66)*), and in the basal chambers by fresh CnT-PR-3D. After incubation for 8 h at 37 °C in 5% CO_2_, the total volume of media in basal and apical chambers was spread onto LB plates for colony forming unit (CFU) counting to confirm the absence of growth after overnight incubation at 37 °C. Inserts were washed twice with PBS and incubated with 1% TritonX-100 for 20 min at 37 °C, 5% CO_2_, and mechanical lysis was performed by scraping the surface and pipetting up and down 10 times. Cell lysates were then serial diluted and spread onto LB agar plates for CFU counting after overnight incubation at 37 °C.

### Recovery and analysis of exfoliated cells

After 3, 6 or 12 hpi, exfoliated host cells in the apical milieu were quantified in 4 and 20 μL samples using Acella 20 and 100 (respectively) sample carriers and the fluidlab R-300 automated cell counter (Anvajo biotech). From the remaining volume, 100 μL was cytocentrifuged at 800 rpm for 5 minutes onto glass slides using a Cytospin 2 centrifuge (Shandon). A hydrophobic pen was used to define the location of cytospun cells for subsequent sample processing. Slides were washed with PBS and fixed overnight with 4% paraformaldehyde (PFA) in PBS (Invitrogen) at 4 °C. The following day, PFA was replaced by 1x PBS and inserts were kept at 4 °C until staining.

### Immunofluorescence staining and microscopy

For the detection of umbrella cell biomarkers, uninfected membranes were washed twice with 1X Hank’s Balanced Salt Solution (HBSS, GIBCO) and stained with 5 µg/ml of wheat germ agglutinin conjugated to Alexa Fluor-555 (WGA555, Invitrogen) in HBSS for 2 hours at RT in the dark. Membranes were then washed with 1X Ca^2+^ and Mg^2+^-free PBS prior to blocking with 5% normal goat serum (NGS, Thermofisher) in 1% PBS/Bovine serum albumin (BSA, Sigma-Aldrich) for one hour. Subsequently, membranes were incubated overnight at 4 °C with primary antibodies diluted in 1% NGS in 1% BSA/PBS at 1:50 for rabbit anti-Cytokeratin-20 (CK20) polyclonal antibody (Invitrogen), or 1:30 for mouse anti-uroplakin-III (UPIII) monoclonal antibody (Invitrogen). After being washed three times in 1% BSA/PBS, membranes were then incubated for 2 hours at RT with a 1:200 dilution of the respective secondary antibody: goat anti-mouse or goat anti-rabbit conjugated to Alexa Fluor-488 (Invitrogen). Membranes were then washed three times in 1% BSA/PBS, and labelling of F-actin was performed with 1:500 dilution of Alexa Fluor-488-conjugated Phalloidin (Invitrogen) for an hour at RT after permeabilization with 0.2% Triton X-100 (Sigma-Aldrich) in PBS for 35 minutes at RT, when WGA was not previously used. DNA were stained with 1 µg/ml 4′′,6-diamidino-2-phenylindole (DAPI, Invitrogen) for 15 min at RT, and membranes were mounted with ProLong™ Glass Antifade Mountant (Invitrogen) onto glass slides for imaging. For infected membranes and slides with cytocentrifuged exfoliated cells, bacteria were labelled with a 1:50 dilution in PBS of either the following primary antibodies at 4 °C overnight: mouse anti-O6 (for CFT073), human anti-O25b (for EC10 and mutant), chicken anti-*P. aeruginosa* (Abcam), rabbit anti-*K. pneumoniae* (Invitrogen) and rabbit anti-*P. mirabilis* (Invitrogen), rabbit anti-LC3A/B (Cell Signaling), rabbit anti-Rab27a (Invitrogen) or mouse anti-UPIIIa (Santa Cruz Biotechnology); or with FITC-conjugated polyclonal antibodies (Invitrogen) at room temperature for 3 h: anti-*Streptococcus* Group B, anti-*E. coli* serotype O/K or anti-*Enterococcus* sp.. Primary antibody incubations were followed by a second incubation with a 1:300 dilution in 1% NGS of goat anti-chicken, anti-rabbit, anti-mouse or anti-human secondary antibodies conjugated with Alexa-Fluor 488 (Invitrogen), respectively, for 2 h at room temperature. WGA or Phalloidin, and DAPI staining and mounting of the samples were performed as aforementioned.

To examine intracellular bacteria by confocal microscopy, the above staining strategies allowed differential colour detection, with extracellular bacteria stained with anti-bacteria FITC-conjugated antibody and blue (DAPI), whereas intracellular bacteria were only stained with blue, and could be distinguished from host cell nuclei by morphology. As a control, the fixation procedure above with and without subsequent membrane permeabilization was used in infected/uninfected organoids that were then stained with a mouse monoclonal anti-mitochondria antibody Cy3 conjugate (Merck) incubated overnight at 4 °C, to confirm that without permeabilization, there was no access of antibodies into the microtissue.

Visualization of stained organoids was performed by confocal laser scanning microscopy using a Leica SP8 microscope, with inspection for intracellular bacteria in top-down views and Z-stacks. Images were processed using the Leica Application Suite (LASX), Advanced Fluorescence 3.1.0 build 8587 Software.

### Scanning electron microscopy (SEM)

Fixation of samples for SEM was performed in 2.5% glutaraldehyde/2% PFA in 0.1 M sodium cacodylate (CAC, TAAB Laboratories) buffer for 30 minutes at RT. Fixed membranes were then incubated with 1% osmium tetroxide (TAAB Laboratories)/1.5% potassium ferricyanide (Sigma-Aldrich) for 1 h at 4 °C, followed by three washes with 0.1 M CAC buffer. Subsequently, they were incubated in 1% tannic acid (TAAB Laboratories) in 0.05 M CAC buffer in the dark at RT for 40 minutes, followed by two washes with 0.05M CAC buffer and one wash with distilled and deionized water (ddH_2_O). Membranes were then dehydrated in graded ethanol (Sigma-Aldrich) series: 2 minutes incubation each in 50%, 70%, 90% and 10 minutes in 100% ethanol, twice. Dehydrated organoids were completely dried with a Leica EM CPD300 critical point dryer. The membranes were then detached from the plastic inserts using a scalpel and were mounted onto aluminium stubs using carbon sticky tabs, where membranes were sputter-coated with 10 nm of gold. SEM images were acquired using a Zeiss Gemini 300 with a working distance of 8 mm, 1.5 kV accelerating voltage using secondary electron (SE2) detector and processed with Zeiss Atlas 5 software. False-colouring of the images was performed using GIMP 2.10.

### Cyto/chemokines profiling of 3D-UHU

The apical milieu from infections at MOI100 after 12 h were centrifuged at 10,000 x*g* for 5 min, followed by another centrifugation at 13,000 x*g* for 8 min, to remove cellular/bacterial debris. Supernatants were used to quantify the amount of cyto/chemokines produced by the urothelial organoids before and after infection using a human premixed multi-analyte customized kit in a Luminex® bead based immunoassay (R&D systems) according to the manufacturer’s instructions. The final suspensions in wash buffer were analysed using a Luminex^®^ 200^TM^ machine, for the analysis of the following 16 analytes: CCL5 (RANTES), CXCL1 (GRO-α), CXCL2 (GRO-β), GM-CSF (CSF2), G-CSF (CSF3), IFN-y, TNF-α, IL1-α, IL1-β, IL6, CXCL8 (IL8), IL18, Lactoferrin, Lipocalin-2 (NGAL), Pentraxin 3 and Osteopontin.

### Statistical analysis

Data were expressed as means ± standard deviation (SD), plotted and analysed using GraphPad Prism version 9.3.1. Statistical significance between replicates was determined using ANOVA followed by Bonferroni’s test.

For quantification of *P. aeruginosa* and *E. coli* rod and filament lengths, confocal images were converted in 16-bit images and analysed in the ImageJ software using the MicrobeJ plugin*(67)*. A total of 1000 bacteria were counted across biological triplicates per strain, using 3-5 regions per replicate, with the exception of COM3 where 15 regions were needed to achieve equivalent total bacterial count. Bacterial lengths were gated from 0-4 μm for rods and >4 μm for filaments, while declustering was manually performed.

## List of Supplementary Materials

fig. S1 to S5

fig. S1. 3D-UHU differentiation marker expression and colonization by *E. coli* clinical isolates and ASB.

fig. S2. Morphology and colonization strategies by non-UPEC uropathogens in the urothelial microenvironment.

fig. S3. UPEC and ASB *E. coli* surface and piliation after infection.

fig. S4. Hemagglutination assays with ASB-like and uropathogenic *E. coli*. fig. S5. 3D-UHU tissue responses to infection.

## Acknowledgments

We would like to thank all funding sources. We thank Pfizer employees Raphael Simon and Robert Donald for review of the manuscript and supplying EC6/10 strains, and David Keeney for construction of FimH knockout strains. We additionally thank Scott Hultgren’s lab for UTI89 and KP. We would also like to thank members of our laboratory for helpful discussion.

## Author contributions

Conceptualization: CF, JLR

Methodology: CF, JF, AL, RGM, AA, IJW, RF, JLR

Investigation: CF, JF, RGM, AA

Funding acquisition: JLR

Project administration: CF, JLR

Writing: CF, JLR

## Data and materials availability

Any additional information or data required to reanalyse the data reported in this paper is available from the corresponding author upon request. Clinical isolates from Royal Free Hospital & UCL, UK will be shared by the lead contact upon request. Clinical isolates and knockout mutants generated from Dept. of Vaccine Design, Immunology & Anti-Infectives, Pfizer, US can be shared upon request via Material Transfer Agreement (MTA).

