## Supplemental Information for "A human urothelial microtissue model reveals shared colonization and survival strategies between uropathogens and asymptomatic bacteria"

**Supplementary Fig. 1. 3D-UHU differentiation marker expression and colonization by *E. coli* clinical isolates and ASB**

**Supplementary Fig. 2. Morphology and colonization strategies by non-UPEC uropathogens in the urothelial microenvironment.**

**Supplementary Fig. 3. UPEC and ASB *E. coli* surface and piliation after infection.**

**Supplementary Fig. 4. Hemagglutination assays with ASB-like and uropathogenic *E. coli.***

**Supplementary Fig. 5. 3D-UHU tissue responses to infection.**

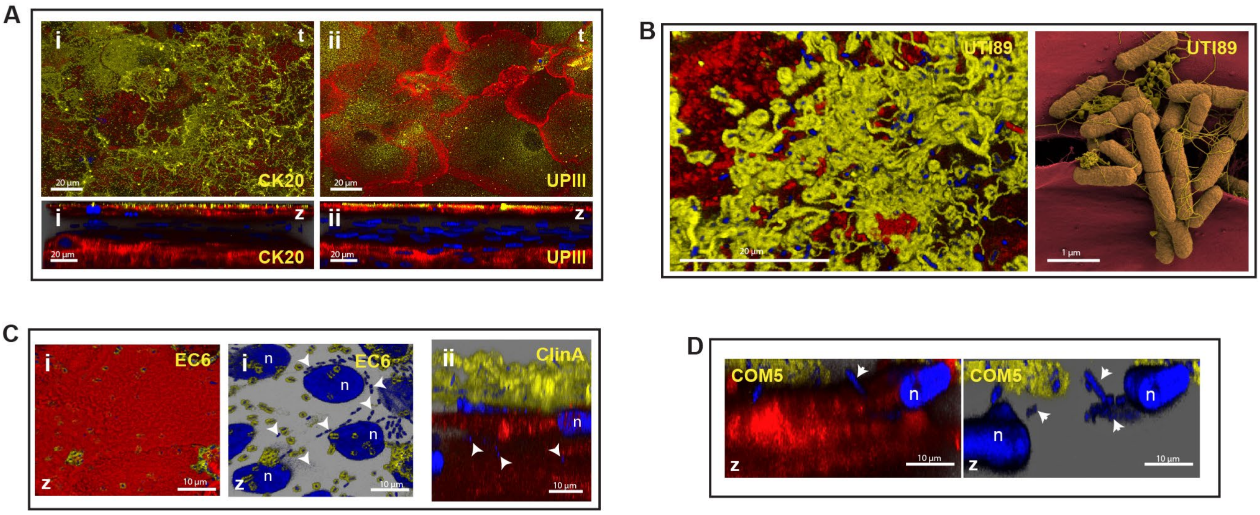


**Supplementary Fig. 1. 3D-UHU differentiation marker expression and colonization by *E. coli* clinical isolates and ASB.** (**A**) Expression of terminal differentiation markers (yellow) cytokeratin 20 (CK20, i) and uroplakin III (UPIII, ii) by apical umbrella cells. (**B**)Biofilm-like aggregates by UTI89. (**C** and **D**) Isolated intracellular bacteria (IIB, arrowheads) by UPEC clinical isolates (C) and ASB COM5 (D). t, top-down view; z, side view. Yellow, extracellular bacteria; Blue, DNA of host nuclei (n) and bacteria (intra and extracellular); Red, cell membrane (CM). Confocal (A,B,C,D) and SEM (B) images representative of a minimum of 4 independent biological replicates per strain.

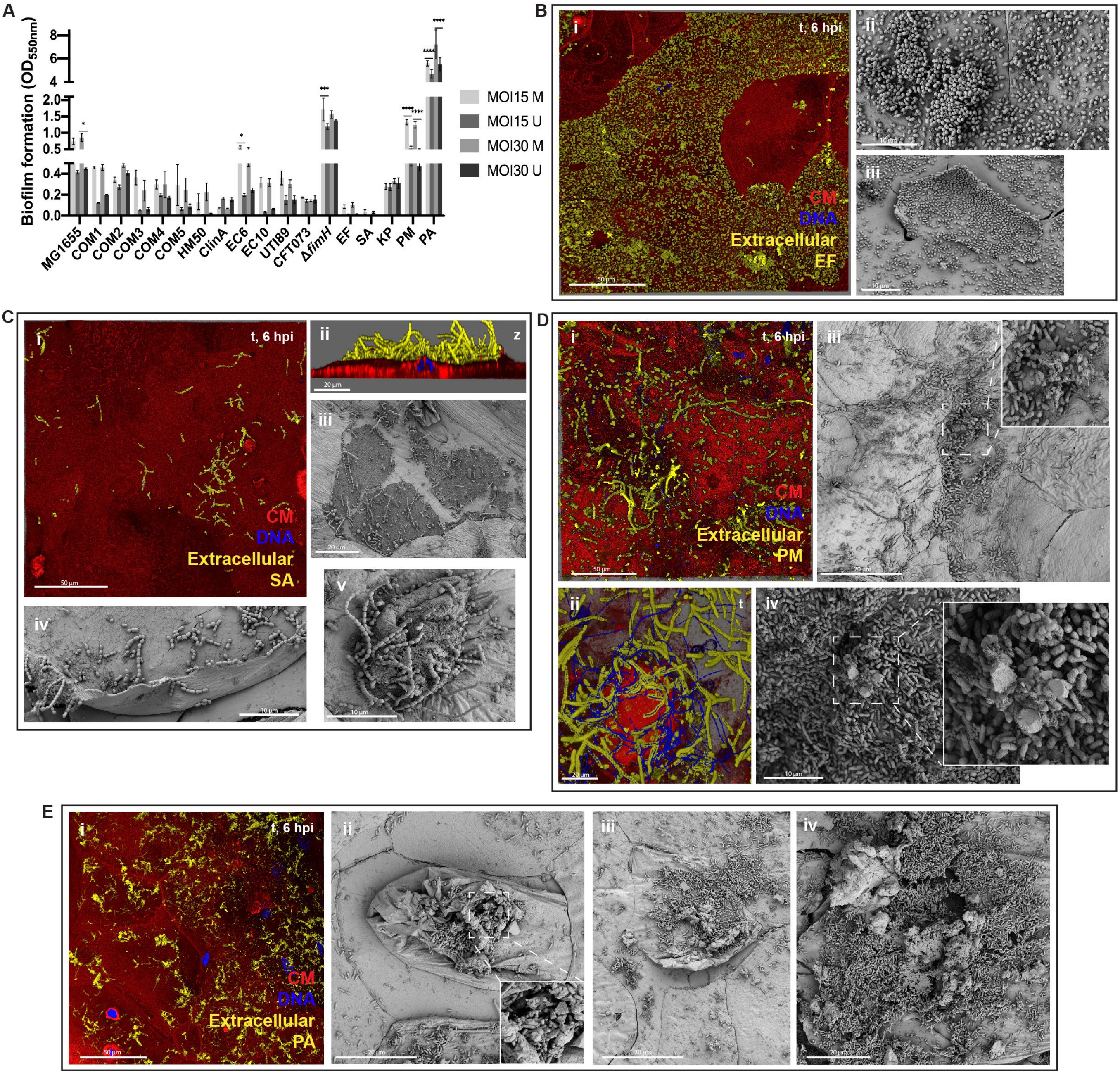


**Supplementary Fig. 2. Morphology and colonization strategies by non-UPEC uropathogens in the urothelial microenvironment.**

(**A**)Assessment of biofilm formation by crystal violet assay using Calgary system after 24 h of bacterial incubation in 25% urine (U) and LB media (M), at MOI 15 and 30. OD at 550 nm is presented as mean ± SE of 2 biological and 4 technical replicates (*, $p < 0.1$; **, $p < 0.01$; ***, $p < 0.001$; ****, $p < 0.0001$). (**B**) *E. faecalis* (EF) spread on the urothelial surface 6 hpi (i), heavily colonized umbrella cells being exfoliated (i) and biofilm-like aggregates (ii). t, top-down view; z, side view. Yellow, extracellular bacteria; Blue, DNA of host nuclei (n) and bacteria; Red, host cell membrane (CM). (**C**) *S. agalactiae* (SA) in discrete regions of the urothelial surface (i-iii), associated with damaged upper cell host membranes (iii). Cocci and chains underneath cell being exfoliated (iv) and dying (v). (**D**) *P. mirabilis* (PM) chains and filamentous forms (i,ii), and interjunctional crystalline biofilm aggregates in the inflamed microtissue (iii,iv). (**E**) *P. aeruginosa* (PA) small biofilm aggregates 6 hpi (i), and larger at 12 hpi with incorporation of exfoliating cell debris and crystalline structures (ii-iv). Confocal (Bi, Ci, Di, Ei) and SEM (Bii,iii, Cii-v, Diii,iv, Eii-iv) images are representative of a minimum of 4 independent biological replicates per strain.

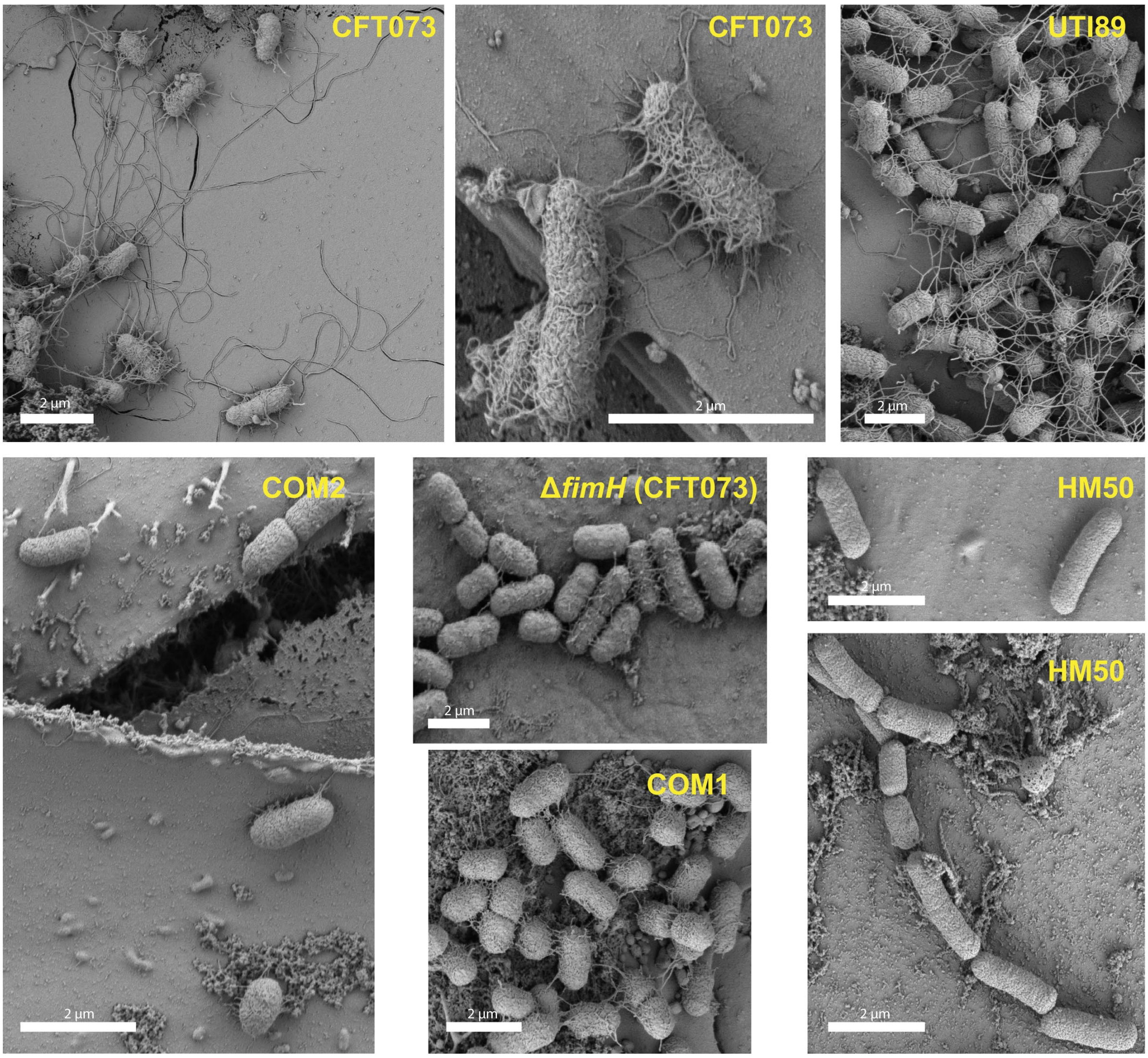


**Supplementary Fig. 3. UPEC and ASB E, coli surface and piliation after infection.** SEM micrographs depicting bacterial surface and piliation of UPEC and ASB-like *E. coli* 12 hpi in the 3D-UHU model. Images are representative of a minimum of 4 independent biological replicates per strain.

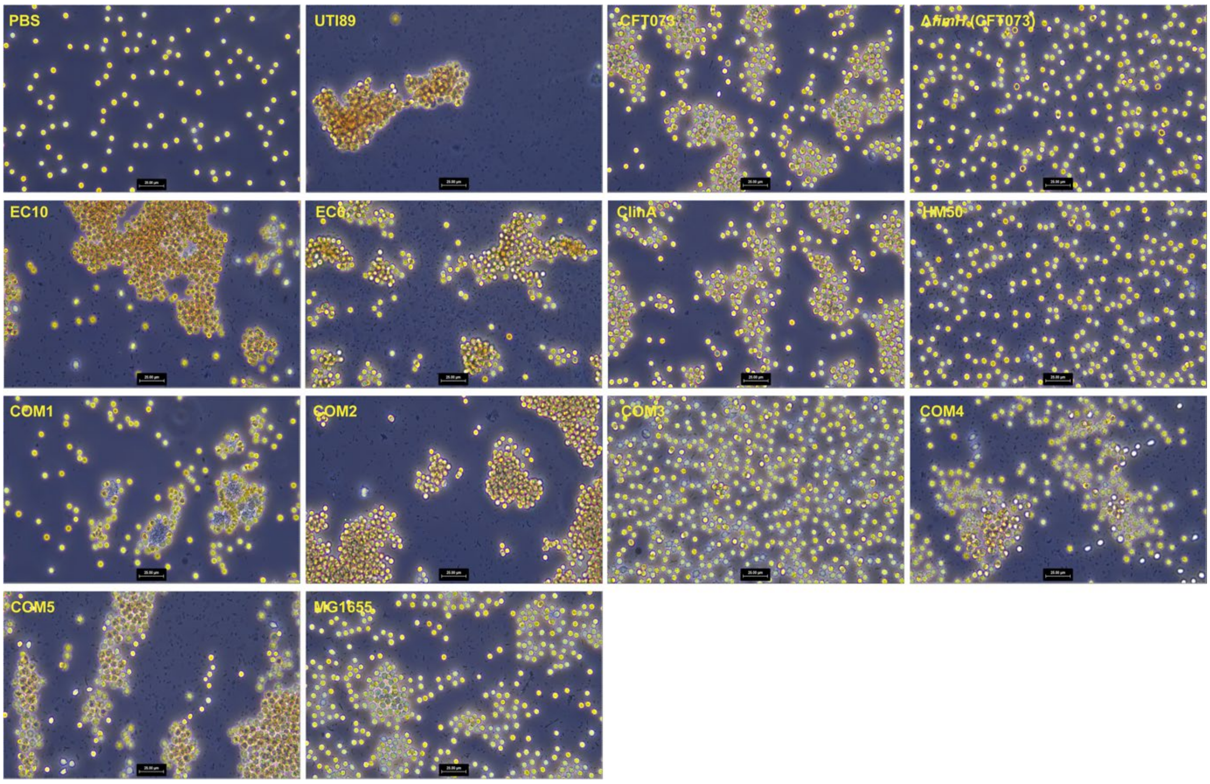


**Supplementary Fig. 4. Hemagglutination assays with ASB-like and uropathogenic *E. coli*.** Brightfield images of hemagglutination of 5% guinea pig red blood cells by ASB and uropathogenic *E. coli* strains after 30 min incubation at room temperature. Scale bars, 25 µm.

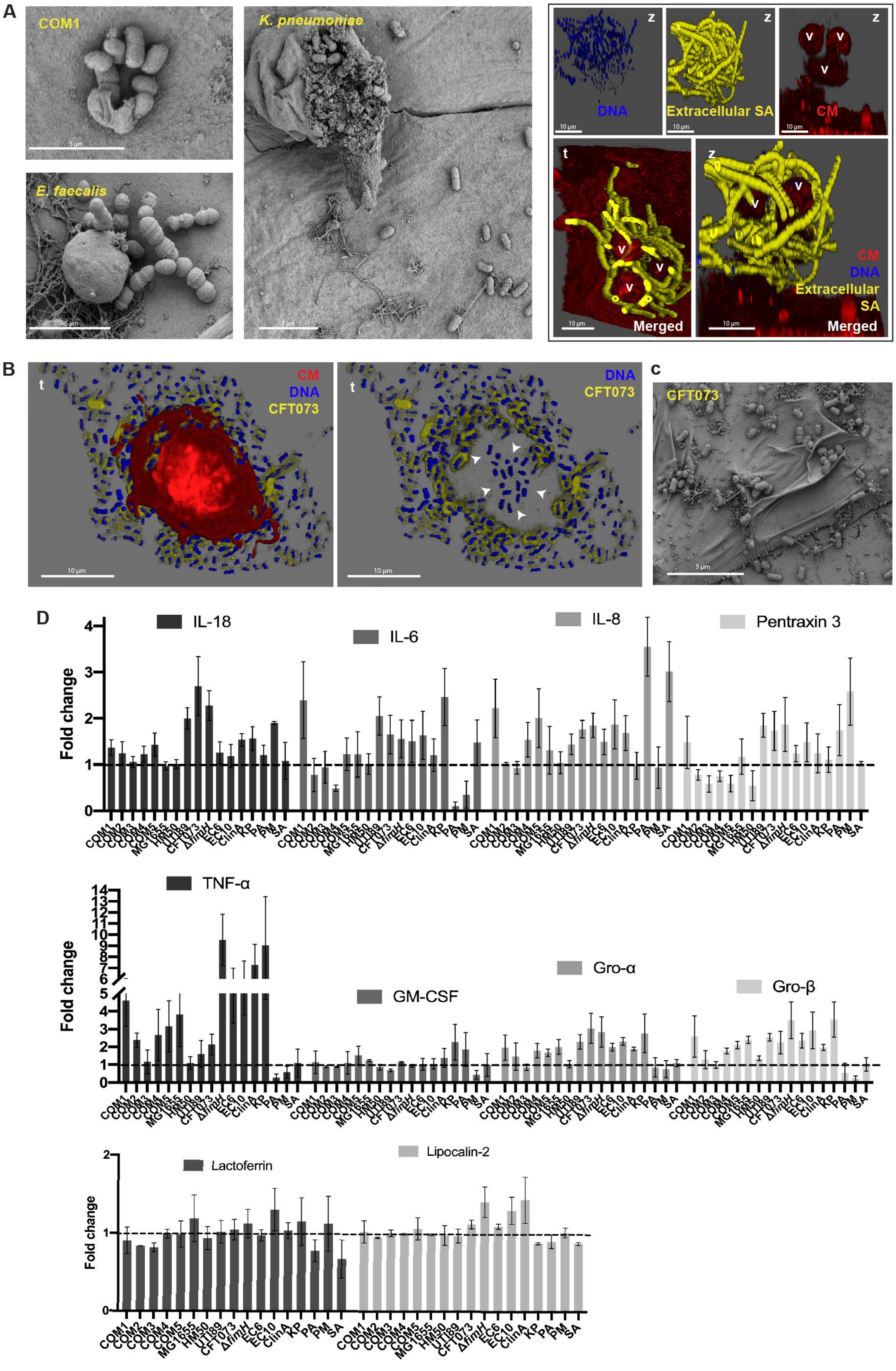


**Supplementary Fig. 5. 3D-UHU tissue responses to infection.**

(**A**) Uropathogens (*E. faecalis*, *K. pneumoniae*) and ASB (COM1) erupting from vesicle-like structures/host cell fragments, and *S. agalactiae* (SA) chains surrounding those structures. Staining as in fig. S2. t, top-down view; z, side view. (**B**) Top-down view of a cytospun cell fragment containing intracellular UPEC CFT073 (arrowheads). (**C**) Umbrella cell membrane ruffling 12 hpi with CFT073. (**D**) Fold change of cyto/chemokine production by 3D-UHU after 12 hpi with ASB-like bacteria and uropathogens. Dashed line represents the mean for uninfected controls, considering three biological and technical replicates. Confocal (SA and b) and SEM (COM1, KP, EF and c) images are representative of a minimum of 4 independent biological replicates per strain.
